# Fluctuation Test with Phenotypic Switching: A Unified Stochastic Approximation Framework

**DOI:** 10.1101/2025.05.20.655090

**Authors:** Anna Hlubinová, Pavol Bokes, Abhyudai Singh

**Affiliations:** Department of Applied Mathematics and Statistics, Comenius University, Bratislava 84248, Slovakia; Department of Electrical and Computer Engineering, Biomedical Engineering, University of Delaware, Newark, DE USA 19716

## Abstract

This paper examines a structurally symmetric fluctuation test experiment in which cell populations grow from a single cell to a set size before undergoing treatment. During growth, cells may acquire tolerance to treatment through probabilistic events, which are passed to progeny. Motivated by recent research on drug tolerance in microbial and cancer cells, the model also allows tolerant cells to revert to a sensitive state, reflecting dynamic phenotypic switching. The master equation governing the probability distribution of tolerant cells is solved via the generating function method and the quasi-powers approximation. Depending on model parameters, the distribution may be approximated by a stable distribution (or its special case, the normal distribution) or through large deviations theory. In the regime of frequent switching, the large deviations approach provides better agreement with numerical solutions, particularly at distribution tails. Conversely, in the regime of infrequent switching, the general stable distributions offer improved accuracy over the Landau distribution, which represents a limiting distribution in case of unidirectional switching.

## I. INTRODUCTION

The Luria-Delbrück fluctuation test demonstrated that the extent of resistance in bacterial populations follows a heavy-tailed distribution, contradicting the Lamarckian hypothesis of induced resistance and supporting the Darwinian view that resistance arises randomly before exposure and is inherited [1], [2]. This discovery led to the development of stochastic models that describe resistance emergence in exponentially growing populations, often modeled using heavy-tailed distributions such as the Lea-Coulson [3], [4], alpha-stable [5], [6], and Fréchet distributions [7], [8].

The classical fluctuation test focuses on resistance driven by genetic heterogeneity. However, isogenic cells can exhibit considerable variability of phenotype due to the random nature of gene expression [9]–[11]. For instance, a small fraction of *E. coli* cells stochastically enter a drug-tolerant persister state, evading antibiotic action [12]–[18] — an effect similarly observed in cancer cells [19]–[23]. Moreover, experiments reveal that individual cells reversibly switch between drug-sensitive and drug-tolerant states driven by the underlying stochastic dynamics of gene regulatory networks [24]–[27]. Phenotypic switching is thought to be reversible, occurring over a range of timescales [28], [29], and impact diverse biological processes, such as stem cells [30]–[32], viral-susceptible cellular states [33]–[35], and metabolic bet hedging [36], [37]. This motivates us to study growing populations with multiple cell states under diverse regimes.

This paper presents a unified framework for deriving approximate distributions in a fluctuation test model with a reversible two-state switching mechanism. Our approach builds upon a quasi-power approximation derived for probabilistic urn models and random tree theory [38]–[40]. By refining this approximation, we systematically develop the classical limit theorem (normal approximation), a large deviation approximation, and a generalized central limit theorem (approximation by alpha-stable distributions). We apply these results to an experimentally relevant case where the initial cell state is drawn from the stationary distribution of the phenotype-switching process.

## II. MODEL FORMULATION

We assume that the population consists of a finite number of cells which can be sensitive or tolerant and that each cell proliferates with a constant intensity which is independent of cell type. Without loss of generality, the value of the proliferation rate can be set to one, i.e. the probability that a cell divides in a time interval (*t, t* + *dt*) is equal to the interval length *dt*.

In order to simplify the modeling, we assume that phenotype switching occurs exclusively during cell division events: a sensitive mother cell has a tolerant daughter cell with probability *µ*, while a tolerant mother cell has a sensitive daughter cell with probability *λ* (Figure 1).

**Fig. 1.**
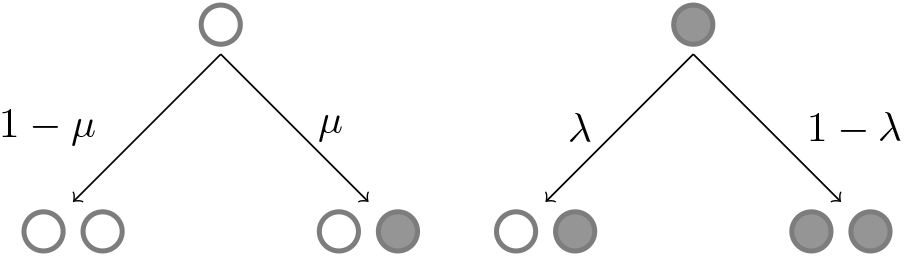
White balls represent sensitive cells, while black balls represent tolerant cells. Arrows indicate the four possible outcomes of cell division: a sensitive cell produces either a sensitive or tolerant offspring with probabilities 1−*µ* and *µ*, respectively, whereas a tolerant cell produces either a sensitive or tolerant offspring with probabilities *λ* and 1 − *λ*, respectively.

The temporal dynamics of the population is modelled by a bivariate Markov process (*m*(*t*), *N* (*t*)), which specifies the total number of cells *N* (*t*) and the number of tolerant cells *m*(*t*) at time *t* (whereby *N* (*t*) − *m*(*t*) gives the number of sensitive cells). The probability that a tolerant cell divides in a time interval (*t, t*+*dt*) is *m*(*t*)*dt* while the probability that a sensitive cell divides is (*N* (*t*) − *m*(*t*))*dt*. Multiplying these by the appropriate phenotype transition probabilities leads the intensities of sensitive/tolerant births in the population process (Table I).

**TABLE 1.** Fluctuation EXPERIMENT MODEL.

| Stochastic event | Changes in population counts | Probability of occurrence in $(t, t + dt]$ |
| --- | --- | --- |
| Sensitive cell birth | $\binom{m}{N} \rightarrow \binom{m}{N+1}$ | $((1 - \mu)(N - m) + \lambda m)dt$ |
| Tolerant cell birth | $\binom{m}{N} \rightarrow \binom{m+1}{N+1}$ | $(\mu(N - m) + (1 - \lambda)m)dt$ |

### A. Difference equation

For a fixed time *t*, the total population size *N* (*t*) exhibits significant variability, which is inconsistent with real-world fluctuation experiments, which are typically evaluated after a certain population size is reached [12]. To address this, we condition the system on reaching a predefined population size *N* and analyze the conditional distribution *P*_*N*_ (*m*), representing the probability of observing *m* tolerant cells in a total population of *N*. This distribution satisfies the following difference equation [41]:

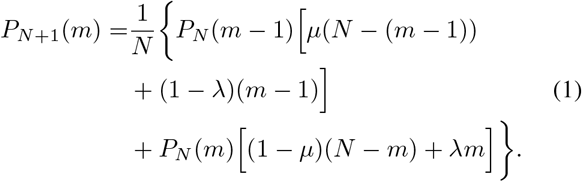

The master equation (1) needs to be complemented by an initial condition which details the composition of the population as it totals *N*_0_ individuals. Typically, *N*_0_ = 1 or a small number. One specific initial condition that we consider is deterministic:

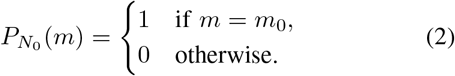

The condition (2) implies that the population starts with *N*_0_ individuals from which *m*_0_ *≤ N*_0_ are tolerant. We will also be focusing on a specific kind of stochastic initial condition

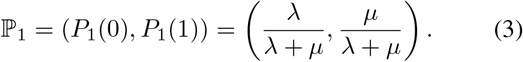

Here, the initial cell is randomly drawn from the steady state distribution of the switching process with forward rate *µ* and backward rate *λ*.

The probability of observing an exclusively sensitive or an exclusively tolerant population can be calculated explicitly

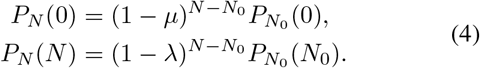

Equations (4) provide boundary conditions for the master equation (1).

### B. Difference–differential equation

Using standard rules [42], we transform the master equation (1) into a difference–differential equation

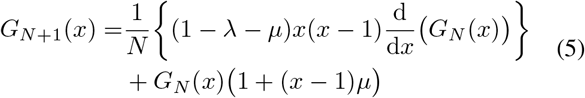

for the probability generating function

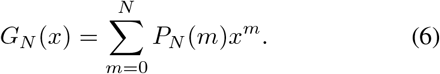

Section III-A is dedicated to deriving a quasi-powers solution to (5). Sections III-B and III-C outline its implications for the asymptotic approximation to the approximate distribution of tolerant cells.

## III. METHODS

In this section, we develop asymptotic approximations to describe the distribution of tolerant cells. The necessary nomenclature is summarized in Table II.

**TABLE 2.**
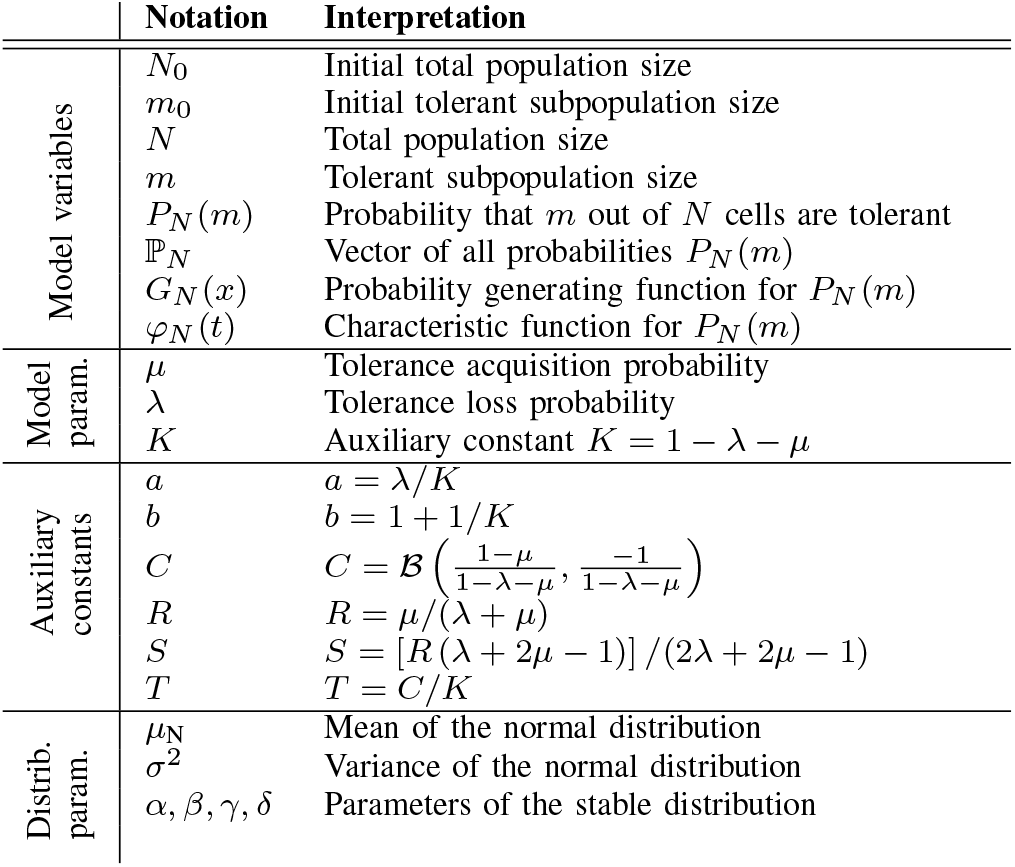
Variables, FUNCTIONS AND CONSTANTS USED IN THIS PAPER.

### A. Quasi-powers ansatz

The fundamental approximation for the generating function (6) is obtained by the quasi-powers ansatz [40]:

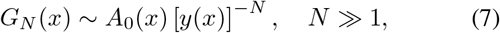

where *A*_0_(*x*) is a pre-exponential factor while *y*(*x*) gives the base of the exponential term.

Inserting the ansatz (7) into (5) and collecting leading-order terms provides the following linear first-order differential equation for the function *y*(*x*):

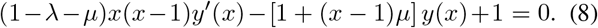

Moreover, it can be shown that *A*_0_(*x*) = *A*_0_ = 1. Thus, it is sufficient to find a function *y*(*x*) that satisfies (8), and

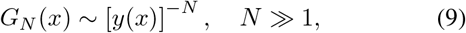

determines the asymptotics of the generating function for large population sizes *N*.

The differential equation (8) has three singular points: *x* = 0, *x* = 1 (in which the term multiplying the derivative vanishes) as well as *x* = *∞*. In the neighborhood of the singular points we require

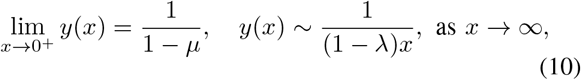

which can be deduced from boundary conditions (4) for the master equation. We also require the continuity at the singular point *x* = 1, specifically that

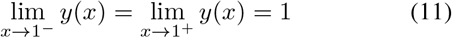

The condition (11) is a direct consequence of the normalization property of the generating function (*G*_*N*_ (1) = 1).

The solution to (8) subject to (10) and (11) is found by the method of separation of variables. For *x ∈* (0, 1) and *K* = 1 − *λ* − *µ >* 0, we find

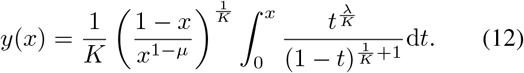

This solution formula will be sufficient for most of our discussion; however, a complete list of formulae that extend to *x >* 1 and *K ≤* 0 are provided in Appendix A.

In Section III-B, we look for the asymptotic expansion of the quasi-powers solution around the interior singular point *x* = 1. This is a prerequisite for the distribution approximations provided in Section III-C.

### B. Asymptotic approximation of y(x) around x = 1

Assume *K >* 0 and *x ∈* (0, 1), representing the asymptotic behavior of the solution (12) as *x →* 1^−^. The case *x →* 1^+^ is discussed in Appendix B.

By setting *δ* = 1 − *x* and *s* = 1 − *t* in (12), the solution as *δ →* 0 is obtained as

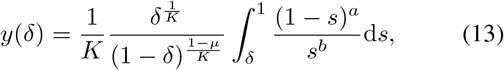

where the notation *a* = *λ/K* and *b* = 1 + 1*/K* was introduced.

The integral *I* appearing in (13) can be evaluated using *Mathematica* software. The result is expressed in terms of the incomplete Beta function, with its expansion given as:

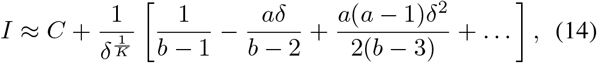

where 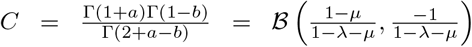 is a constant.

The substitution of the expansion (14) into (13), along with the application of the Taylor series expansion to the expression

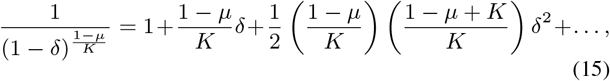

as *δ→* 0, followed by back-substitution into the variable *x*, yields the result:

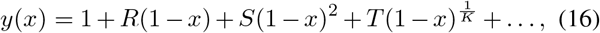

where *R* = *µ/*(*λ* + *µ*), *S* = [*R* (*λ* + 2*µ* − 1)] */*(2*λ* + 2*µ* − 1) and *T* = *C/K*.

It is important to note that, in addition to the regular powers (1−*x*)^0^, (1−*x*)^1^, (1−*x*)^2^, … the expansion (16) also includes the non-integer powers 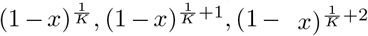, …. For the purposes of this analysis, it is sufficient to consider the three largest terms in the expansion. Depending on the value of *K*, the second-order term would either be (1 − *x*)^2^ (for 0 *< K <* 1*/*2) or 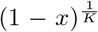 (for 1*/*2 *< K <* 1). Moreover, it is important to observe that the approximation (16) remains valid up to second order for *K <* 0, with deviations occurring exclusively in higher-order terms.

### C. Theoretical approximations of P_N_ (m)

The aim of this section is to derive three approximations of the probability distribution *P*_*N*_ (*m*) depending on the value of *K*. Without loss of generality, consider the asymptotic expansion (16) for *x ∈* (0, 1).

1. *Normal distribution:* As demonstrated above, for 0 *< K <* 1*/*2, the second-order term in the approximation of the function *y*(*x*) around *x* = 1 is the regular second power, (1 − *x*)^2^. It is therefore hypothesized that for 0 *< K <* ½ a reasonable approximation of the probability distribution of *P*_*N*_ (*m*) is the normal distribution with mean *µ*_N_ and variance *σ*^2^. (The subscript *N* at the mean is introduced to distinguish it from the mutation coefficient, denoted by *µ*.) By comparing the characteristic function *φ*_*m*_(*t*) of *P*_*N*_ (*m*)

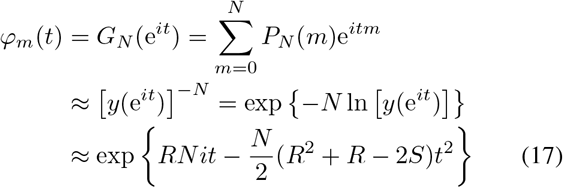

with the characteristic function of the normal distribution

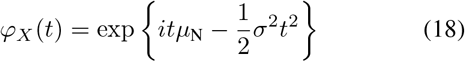

the parameters of the normal distribution

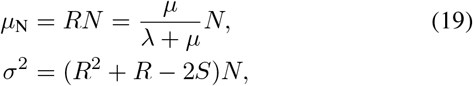

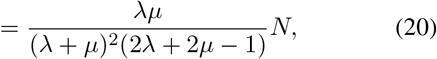

are obtained. It is evident that for the dispersion (20) to be a positive number, the condition *K <* 1*/*2 must be met. The parameters obtained for the normal distribution are consistent with the result of Example 7.3 (Randomized play-the-winner) from [43].
2. *Stable distribution:* If 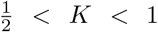, the second-order expansion includes a term with an anomalous exponent 1*/K*. In this regime, the probability distribution *P*_*N*_ (*m*) is expected to align closely with a class of stable distributions. Results in [41] show that for small *µ* and *λ* (*∼* 10^−3^), *P*_*N*_ (*m*) is well-approximated by the Landau distribution, parametrized in [44] as 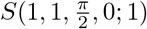. By comparing the characteristic function *φ*_*m*_(*t*) of *P*_*N*_ (*m*)

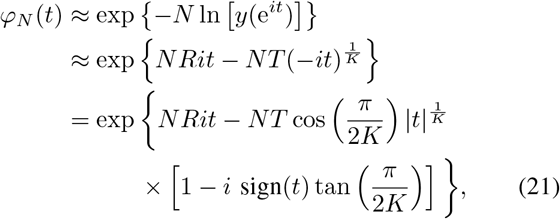

with the characteristic function *φ*_*X*_ (*t*) of stable distribution parametrized as **S**(*α, β, γ, δ*; 1)

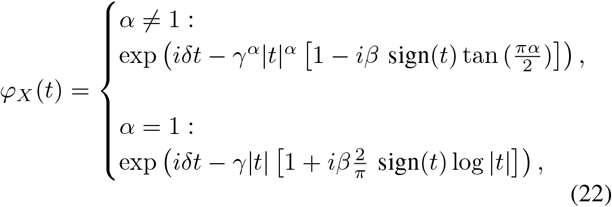

parameters of the stable distribution

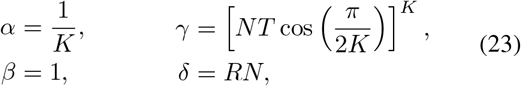

are obtained. It is important to note that the Landau distribution 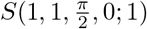 represents a limiting case of the more general class of stable distributions. Additionally, the normal distribution is a specific case within the broader stable distribution family.
3. *Theory of large deviations:* The theory of large deviations examines the exponential decay of probabilities associated with large fluctuations in random systems [45]. This work demonstrates the application of this framework to approximate the probability distribution in the Luria–Delbrück model.

Let *A*_*N*_ be a real-valued random variable parameterized by a positive integer *N*, representing the fraction of tolerant cells in a population of size *N*. The realizations of *A*_*N*_ take values *a* = *m/N ∈* (0, 1). The cornerstone of large deviation theory is the *large deviation principle*, which establishes an exponential approximation for probabilities:

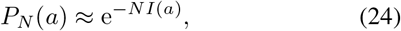

where *N* is typically a large parameter and *I*(*a*) is a continuous function known as the *rate function*.

A key result within this theory is the Gärtner–Ellis theorem [45], which provides conditions under which the large deviation principle holds. Specifically, if the *scaled cumulant generating function* of *A*_*N*_, defined as

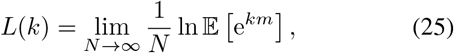

exists and is differentiable for all *k ∈* ℝ, then *A*_*N*_ satisfies the large deviations principle in (24). The corresponding rate function *I*(*a*) is then determined via the Legendre–Fenchel transformation of *L*(*k*) as

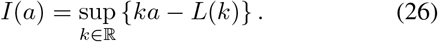

By utilizing the approximation (9) for the generating function and the relationship between the generating function and the expected value, expressed as

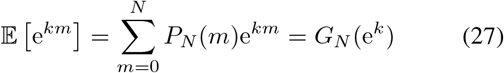

the function *L*(*k*) can be approximated as

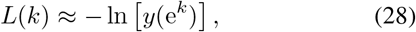

where *y*(*x*) is determined by the solutions derived above. Given the definition of the function *L*(*k*), its differentiability for *k ∈* ℝ is equivalent to the differentiability of the function *y*(*x*) for *x >* 0, which is satisfied.

## IV. RESULTS

This section compares the approximations of probability distribution *P*_*N*_ (*m*) from the previous section with numerical results, using graphical analysis to assess accuracy across varying *µ* and *λ*. Three cases are distinguished based on the value of *K* = 1 = *λ* − *µ*. The value of *K* lies in the interval (−1, 1) and characterizes the switching intensity. If *K* = −1, then the daughter cell deterministically adopts the opposite type. If *K* = 0, the daughter cell’s type is chosen at random. If *K* = 1, then the daughter cell always adopts the cell type of the mother.

### A. Case −1 < K < 0

Figure 2 compares the numerical solution with the normal distribution approximation and the large deviations approximation for the symmetric initial conditions and unequal *µ* and *λ*. The near overlap of all three curves in Figure 2 confirms the excellent performance of both approximations in this case.

**Fig. 2.**
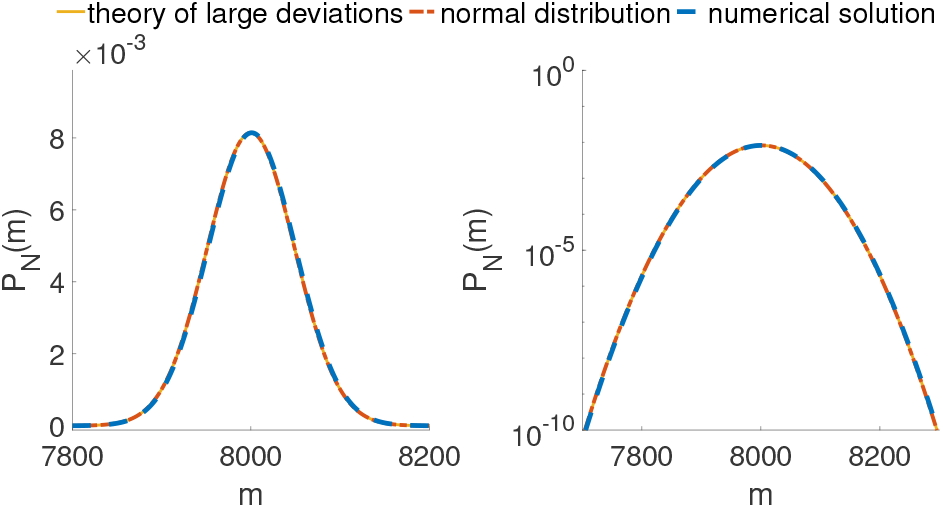
Comparison of the probability vector ℙ_*N*_ obtained numerically, the normal distribution density with mean (19) and variance (20), and the approximation using the large deviation theory. Parameters: 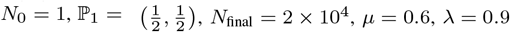, i.e., *K* = −0.5.

### B. Case 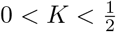

In the previous section, the approximation using the normal distribution for values of *K* within the specified range was derived, alongside an alternative approximation based on the theory of large deviations. Figure 3 compares these two approximations with the numerical solution under symmetric initial conditions and unequal parameters *µ* and *λ*. The logarithmic scale plot of *P*_*N*_ (*m*) highlights a marked difference between the (by definition) symmetric normal distribution approximation and the asymmetric approximation obtained via the theory of large deviations. Notably, the latter almost perfectly aligns with the numerical solution, underscoring its significant contribution.

**Fig. 3.**
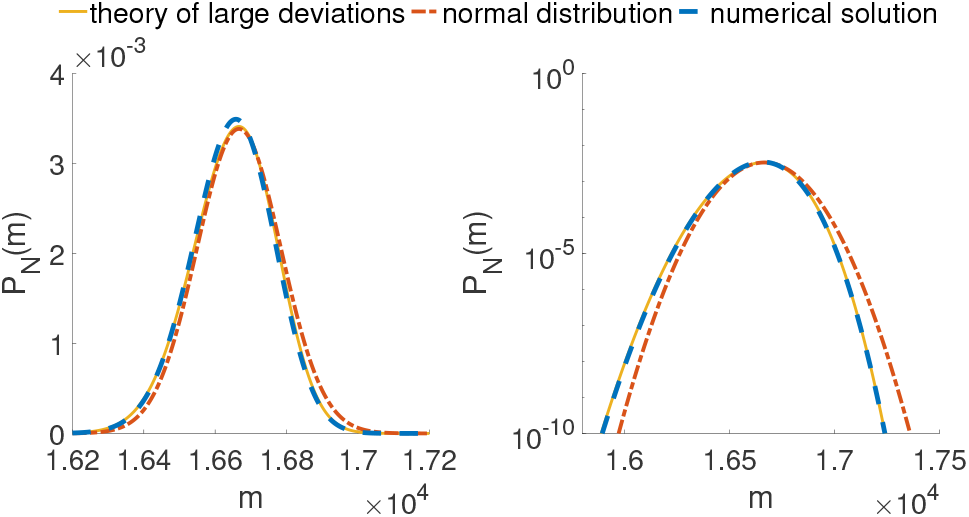
Comparison of the probability vector ℙ_*N*_ obtained numerically, the normal distribution density with mean (19) and variance (20), and the approximation using the large deviation theory. Parameters: 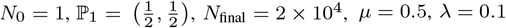, i.e., *K* = 0.4.

### C. Case 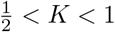

In this case, the second-order expansion of the solution *y*(*x*) around *x* = 1 includes a term with an anomalous exponent 1*/K*. The normal distribution approximation fails here. However, a general stable distribution with parameters derived in (23) is expected to provide a valid alternative. The accuracy of this approach is compared with the Landau distribution [41]. Moreover, as Figure 4 shows, for *K* values close to 1*/*2, the large deviations approximation of the probability distribution *P*_*N*_ (*m*) remains highly effective.

**Fig. 4.**
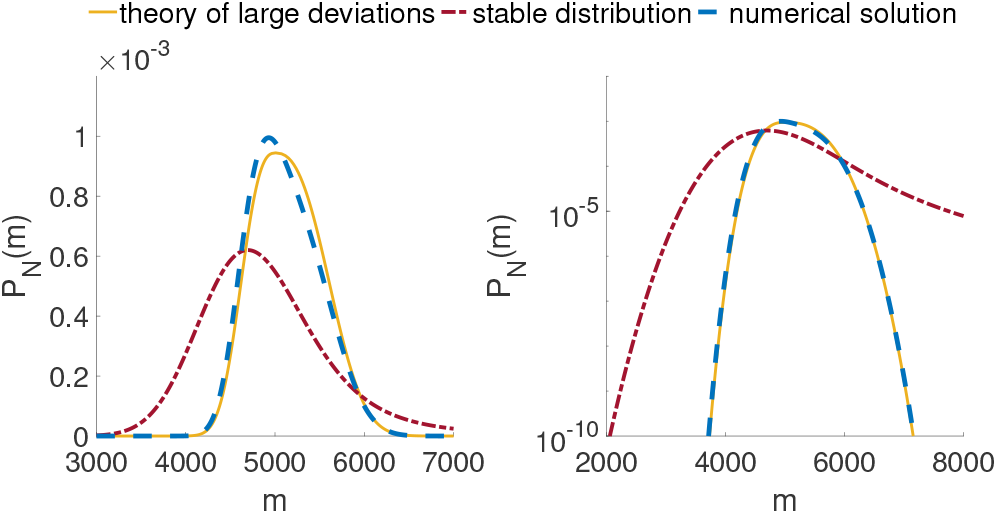
Comparison of the probability vector ℙ_N_ obtained numerically, the stable distribution density with parameters (23), and the approximation using the large deviation theory. Parameters: 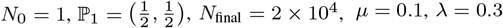 i.e., K = 0.6.

Figure 4 indicates that the theory of large deviations performs exceptionally well even in the case of *K→* 1*/*2^+^ under symmetric initial condition even with unequal parameters *µ* and *λ*. Conversely, the stable distribution fails to provide an accurate approximation of the numerical solution. Two potential explanations for this discrepancy are proposed. First, for small values of the perturbation parameters *µ* and *λ*, the final numerical solution is significantly influenced by the choice of initial conditions. Second, for *K* near 1*/*2, the expansion of the solution (16) becomes increasingly complex due to the importance of higher-order terms.

The numerical solution for *K→* 1 exhibits a strong dependence on the initial condition. In contrast, the approximation based on the theory of large deviations is independent of the initial condition, which accounts for its failure as *K →* 1. As illustrated in Figure 5, the stable distribution with parameters (23) performs well under an asymmetric initial condition for both *K* = 0.89 (left) and *K* = 0.98 (right). Consequently, it provides a more general approximation compared to the Landau distribution, which has been used in the unidirectional case [41].

**Fig. 5.**
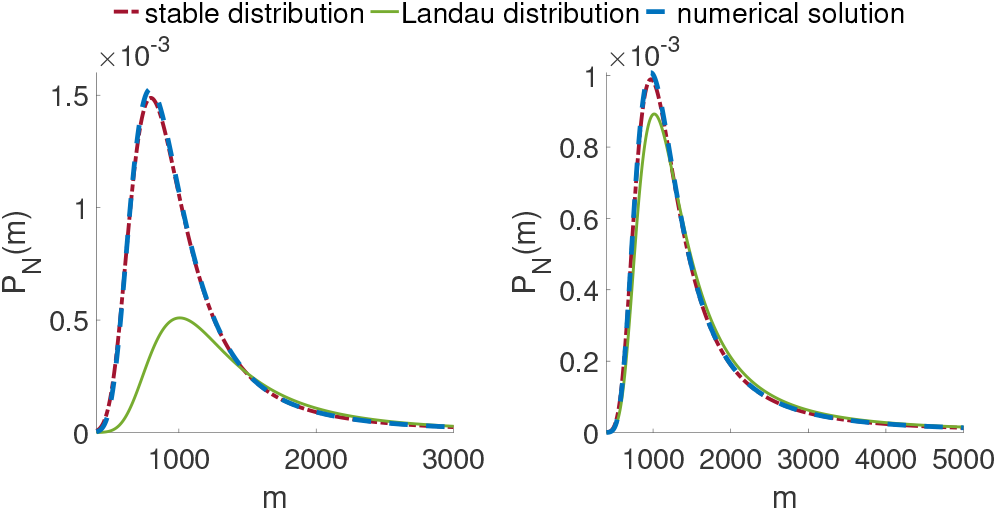
Comparison of the probability vector ℙ_*N*_ obtained numerically, the stable distribution density with parameters (23), and the Landau approximation from [41]. Parameters: *N*_0_ = 1, ℙ_1_ = (1, 0), *N*_final_ = 2 *×* 10^4^. Left: *µ* = 0.01, *λ* = 0.1, i.e., *K* = 0.89. Right: *µ* = 0.01, *λ* = 0.01, i.e., *K* = 0.98.

## V CONCLUSIONS

In this paper, we presented a unified methodology based on a quasi-powers expansion for the construction of approximate distributions in a stochastic fluctuation test model. The normal and large deviation approximations are suitable for fast to moderate switching regimes (Figures 2, 3, and 4) while the stable distribution approximation is suitable for slow switching regimes (Figure 5).

The model is structurally symmetric, from which it follows that the transformation *λ* ⇆ *µ* and *m* ⇆ *N* − *m* provides a new solution to the model and also potentially yields a new approximation. While the normal approximation is invariant under this transformation, for the stable approximation the transformation leads to a reflection around the *m* = *N/*2 axis. Further approximations can be constructed as a convex combination of the original stable distribution and its reflection.

The reflection procedure can be applied to obtain approximations for general initial conditions. Specifically, the type of the initial cell randomly drawn from the steady state switching distribution, i.e. a tolerant cell with probability *µ/*(*λ* + *µ*) or a sensitive cell with the complementary probability *λ/*(*µ* + *λ*). The same steady state values can be achieved by either large or small switching probabilities. The frequent switching case leads to a narrow normal distribution of tolerant cells around the expected value (Figure 6, top). The infrequent switching case, on the other hand, leads to a mixture of the skewed stable distribution and its reflection (Figure 6, bottom). The current unified approximation framework thus provides suitable approximations both for unimodal distributions in the frequent switching case as well as bimodal U-shaped distributions in the case of infrequent switching case.

**Fig. 6.**
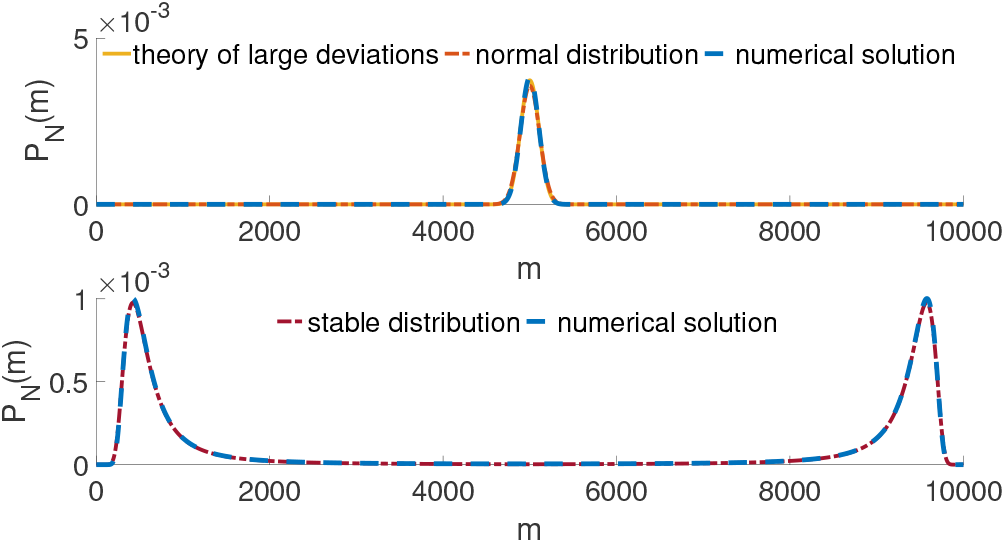
Parameters: 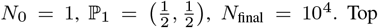: fast-switching, *µ* = 0.3, *λ* = 0.3, i.e., *K* = 0.6. Bottom: slow-switching, *µ* = 0.01, *λ* = 0.01, i.e., *K* = 0.98. *Note: The y-axis scales differ between the upper and lower figures*.

In conclusion, this study provides an approximation framework for a structurally symmetric model of a fluctuation test experiment. The approximations provide a nuanced understanding of Markovian switching in a growing population for both frequent and sporadic switching regimes. Future work should include the effects of additional factors such as environmental changes and multiplicity of cell types.

## Appendix A

### Solution for *y*(*x*)

- *Case K* = 0: If *K* = 1 − *λ* − *µ* = 0, the differential equation (5) reduces to the algebraic equation with a simple solution

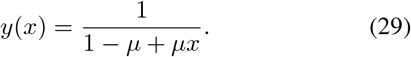 Note that in this case the phenotype of the offspring is independent of the phenotype of the parent. Therefore, the distribution of tolerant cells follows a binomial distribution describing *N* independent experiments, each with the success probability *µ*. The generating function of the binomial distribution is then given by *G*_*N*_ (*x*) = (1 − *µ* + *µx*)^*N*^, where each experiment corresponds to the birth of a cell, and success represents the new cell being tolerant. Thus, in this special case, equality holds in the relation (9), with the solution *y*(*x*) given by (29).
- *Case K*≠ 0: If *K* ≠ 0, the first-order linear differential equation yields a general solution for *y*(*x*) as:

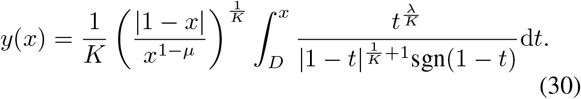 It is necessary to determine the constant *D* that appears in the lower bound of the integral in the solution separately for the domain *x ∈* (0, 1) and for the domain *x >* 1. Moreover, the value of *D* depends on the sign of the constant *K*. The exact solutions are:

- Case *x ∈* (0, 1), *K >* 0: the solution (12).
- Case *x ∈* (0, 1), *K <* 0:

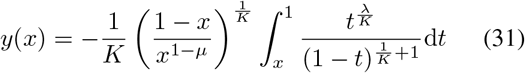
- Case *x >* 1, K > 0:

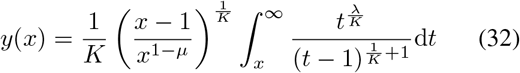
- Case *x >* 1, K > 0:

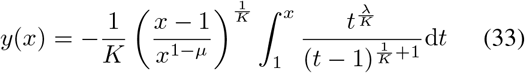

### Appendix B Asymptotic approximation

The asymptotic expansion for the case *x →* 1^+^ is derived from the solution (32) in a manner analogous to the procedure applied for *x →* 1^−^ in Section III-B. This yields the following expression:

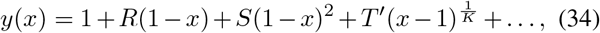

where *R* and *S* retain their previous definitions, *T* ^*′*^ = *C*^*′*^*/K* and 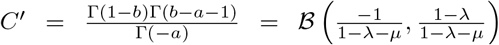 is a constant.

It is worth noting once again that, in the case of *K <* 0, discrepancies with the approximation (34) emerge solely in terms of order higher than second.

## Notes

* This work was supported by the Slovak Research and Development Agency under the contract No. APVV-23-0039 and the VEGA grant 1/0264/25.

### Competing Interest Statement

The authors have declared no competing interest.

